# Midfacial retrusion and loss of facial appendages caused by mutation of Pax9 in zebrafish

**DOI:** 10.1101/2024.12.12.628204

**Authors:** Sandhya Paudel, Sarah McLeod, Stefani Gjorcheska, Lindsey Barske

## Abstract

Loss of dentition has occurred repeatedly throughout vertebrate evolution. Cyprinid fish, including zebrafish, form teeth only deep within the pharynx, not on their oral jaws. However, zebrafish still robustly express transcription factors associated with mammalian tooth development in the neural crest-derived mesenchyme surrounding the mouth. We investigated whether this expression is vestigial or whether these factors contribute to the formation of non-tooth mesenchymal structures in the oral region, using Pax9 as a test case. Zebrafish homozygous for two different *pax9* mutant alleles develop the normal complement of pharyngeal teeth but fail to form the premaxilla bone, most of the maxilla, and nasal and maxillary barbels. Lack of most of the upper jaw complex does not preclude effective feeding in the laboratory environment. We observed a significant deficit of *sp7*:EGFP^+^ osteoblasts and adjacent *alx4a*:DsRed^+^ condensing mesenchyme where the maxilla forms, and no accumulation of either in the premaxillary domain. These phenotypes are not presaged by major disruptions in early facial patterning, Wnt signaling, proliferation, or cell death; however, loss of a small population of Wnt-responsive cells around the maxilla correlates with its stalled growth in mutants. We conclude that Pax9 is not unequivocally required for all vertebrate tooth development, but instead may be broadly involved in the development of a variety of organs forming through mesenchymal condensation around the mouth.

## INTRODUCTION

More than sixty heterozygous variants leading to deletion, nonsense, frameshift, or missense mutations in the transcription factor *PAX9* have been associated with non-syndromic autosomal dominant oligodontia (OMIM #604625: Tooth Agenesis, Selective 3) (Stockton et al., 2000; Nieminen et al., 2001; Das et al., 2002; Frazier-Bowers et al., 2002; Lammi et al., 2003; Chu et al., 2023). Oligodontia is defined as a congenital absence of six or more permanent teeth. In one extreme case, a patient with a R47W missense variant in the DNA-binding domain was missing a total of 20 teeth: four maxillary premolars, three canines, nine molars, and all central incisors (Zhao et al., 2007). Homozygous *Pax9^lacZ/lacZ^*mouse mutants lack erupted teeth entirely: they form tooth buds, but condensation of mesenchymal cells around the bud is reduced, and they fail to progress to cap stage (Peters et al., 1998). Primary tooth induction signals, including *Wnt10a/b*, *Shh*, *Fgf8*, and *Pitx2*, originate in the overlying odontogenic oral epithelium (reviewed by Thesleff et al., 1995; Hermans et al., 2021). Fgf8 induces *Pax9* expression in the underlying neural crest-derived odontogenic mesenchyme (Neubuser et al., 1997), where it in turn drives tooth progression through targets including *Bmp4*, *Msx1*, and *Osr2* (Ogawa et al., 2006; Zhou et al., 2011).

Pax transcription factors are classified by their highly conserved N-terminal Paired box DNA-binding domains, which in some family members co-occurs with a C-terminal homeodomain and conserved octapeptide sequence. Pax9 is most closely related to Pax1, with both possessing the octapeptide but lacking the homeodomain. Pax1 is not involved in mammalian tooth development but collaborates with Pax9 in development of the vertebral column (Peters et al., 1999; Rodrigo et al., 2003). Pax9 is also involved in closure of the secondary palate, patterning of the distal limbs, and formation of organs derived from the pharyngeal endoderm (thymus, parathyroid, and ultimobranchial bodies) (Peters et al., 1998; Zhou et al., 2013).

Despite tooth loss being the most sensitive phenotype in humans carrying heterozygous *PAX9* variants, evidence that Pax9 is not sufficient for tooth development comes from the fact that it is still robustly expressed underneath the oral epithelium in cypriniform fishes (Stock et al., 2006), which lost their oral dentition in a common ancestor 50-100 million years ago (Jackman et al., 2024). This group includes the zebrafish (*Danio rerio*), which form teeth only deep in the pharynx on the fifth ceratobranchials (Huysseune et al., 1998). Other fish species commonly studied in the lab, e.g. medaka (*Oryzias latipes*), three-spine stickleback (*Gasterosteus aculeatus*), Mexican tetra (*Astyanax mexicanus*), and Lake Malawi cichlid spp., are not cyprinids and have oral teeth on their upper and lower jaws. While some putative tooth-inducing epithelial genes like *dlx2a/b* are present in the oral epithelium of tooth-bearing fish but missing in zebrafish (Stock et al., 2006), others are still expressed, including *pax9*’s activator *fgf8a* as well as *shha*, *bmp4*, and *pitx2c* (Balczerski et al., 2012). Odontogenic mesenchyme markers like *barx1*, *msx1a*, *osr1/2*, and *lhx6a* are also still expressed in the underlying oral mesenchyme with *pax9*, highly enriched around the lateral corners of the forming mouth during early facial patterning stages (Stock et al., 2006; Swartz et al., 2011). Recent work has suggested that reduced availability of retinoic acid in the cyprinid face could be the key upstream difference explaining their lack of oral tooth initiation (Jackman et al., 2024). Indeed, the gene encoding the RA-metabolizing enzyme CYP26B1 is highly enriched in zebrafish frontonasal crest (Mitchell et al., 2021). We were interested in whether the continued expression of odontogenic mesenchyme markers in cyprinids is vestigial or if these transcription factors are involved in the development of other non-tooth, neural crest-derived structures around the mouth.

We previously performed short-term lineage-tracing in zebrafish for one of these mesenchymal genes, *barx1*, using a Gal4ff knock-in line, observing that some *barx1*-expressing cells around the mouth are incorporated into the precartilaginous condensations that develop into the larval upper and lower jaw cartilages (Paudel et al., 2022). Functionally, Barx1 suppresses formation of ectopic joins within the lower jaw (Meckel’s cartilage) and drives elongation of the pterygoid process of the palatoquadrate (Nichols et al., 2013). This structure connects the cartilaginous jaw skeleton to the palate and provides the template for the palatine bone, which in turn articulates with the intramembranous maxilla bone (Cubbage and Mabee, 1996). Other *barx1^Gal4ff^*^+^ cells in the oral region perdured as unidentified mesenchyme loosely affiliated with the developing skeleton at the end point of our analysis (Paudel et al., 2022). Some of these cells may contribute to the later-forming maxillary and dentary bones (Cubbage and Mabee, 1996), which were small and dysmorphic in zebrafish *barx1* mutant larvae at 6 dpf (Nichols et al., 2013). The precise developmental origins of these bones in fish have not been mapped. Supporting conserved involvement of *barx1* in tooth development, pharyngeal teeth formed but were detached from their supporting bone, the fifth ceratobranchial, in *barx1* mutants (Nichols et al., 2013).

As for the other odontogenic mesenchyme markers, craniofacial phenotypes for zebrafish *msx1a*, *osr1*, *osr2*, and *lhx6a* null mutants have not yet been reported. A *pax9* morphant zebrafish model presented abnormalities in the dorsal hyoid cartilage and palate as well as reduction of pharyngeal teeth number, but the upper and lower jaw cartilages were normal, and other mouth-adjacent structures forming at later stages were not investigated (Swartz et al., 2011). A report focused on the requirement of Pax9 in granulopoiesis noted jaw defects later in development, but did not further elaborate (Pak et al., 2021).

In this study, we used two new *pax9* mutant models to directly test whether a functional requirement for Pax9 persists in the oral region of zebrafish despite evolutionary loss of teeth. Our results indicate that non-tooth oral structures originating in neural crest-derived mesenchyme are truncated or fail to form entirely in the absence of Pax9. We posit that Pax9’s ancestral role in the oral region may have been to support the development of a variety of organs forming through epithelial-mesenchymal crosstalk and/or mesenchymal condensation, with its activity growing progressively more restricted to the oral teeth in the lineage leading to mammals.

## MATERIALS & METHODS

All zebrafish (Danio rerio) were maintained and handled in accordance with best animal practices as defined by national and local animal welfare agencies. All animal experiments performed in this study were approved by the Institutional Animal Care and Use Committees of Cincinnati Children’s Hospital Medical Center (No. IACUC2024-0108) and the University of Southern California (No. 10885).

### Zebrafish husbandry and published lines

Zebrafish (*Danio rerio*) embryos were raised at 28.5°C in Embryo Medium (EM) from day 0 till 4-6 days post fertilization (dpf) following standard procedures (Westerfield, 2007) and staged as described (Kimmel et al., 1995; Parichy et al., 2009). EM was changed once per day, and any unfertilized or dying embryos were removed. At 4-6 dpf, after the swim bladder had inflated, 20-25 larva were placed in a 3-liter tank and raised in the aquatics facility to 14 dpf on a rotifer or Gemma (Skretting) diet in a gradually increasing water volume. Continuous flow of system water was activated at 14 dpf, and brine shrimp and flake food were gradually added to the diet.

Published mutant and transgenic lines used in this study include *barx1^Gal4ff-ci3030^*(Paudel et al., 2022), *Tg(UAS:nlsEGFP)^el609^* (Barske et al., 2020), *Tg(sp7*:*EGFP*)*^b1212^* (DeLaurier et al., 2010), *Tg(scxa:mCherry)^fb301^* (McGurk et al., 2017), *Tg(Hsa.RUNX2:mCherry*)*^zf3244^*(alias *RUNX2:mCherry*) (Barske et al., 2020), *Tg(fli1:EGFP)^y1^*(Lawson and Weinstein, 2002), *TgBAC(alx4a:DsRed2)^pd52^* (alias *alx4a:DsRed*) (Nachtrab et al., 2013), *Tg(Mmu.Sox10-Mmu-Fos:Cre)^zf384^* (alias *SOX10*:*Cre*) (Kague et al., 2012), *Tg(actb2:LOXP-BFP-LOXP-DsRed)^sd27^*(alias *actb2:BFP>DsRed*) (Kobayashi et al., 2014), and *Tg(7xTCF-Xla.Sia:GFP)^ia4^*(alias *TCF:EGFP*) (Moro et al., 2012). Lines were maintained and used as hemizygotes. Larvae were sorted at 2-6 dpf for carriers of each fluorescent marker. *SOX10:Cre* does not have a linked marker and was maintained by sorting for DsRed expression on the *actb2:BFP>DsRed* background.

### Generation of pax9 mutant alleles and genotyping

The *pax9^el622^* zebrafish mutant line (NM_131298.1:c.243_252del) was generated via TALEN-based targeted mutagenesis as described (Sanjana et al., 2012; Barske et al., 2016). TALENs were designed to target 5’-TCCGGCTCAGGATAGTAGAG-3’ and 5’-TGCTGATGTCACAAGGCCTG-3’, both in the second coding exon. In the *el622* isolate, a 10-bp deletion between these target sites destroys a BsrI site and causes a frameshift and premature termination of translation after 16 incorrect amino acids. *pax9^el622^* fish were genotyped using primers F3 (5’-TCGGAACAGGTCAGAATAGGA-3’) and R3 (5’-TCGGAACAGGTCAGAATAGGA-3’) and Bsrl digestion (New England Biolabs), producing digested wild-type bands (611 and 186 bp) and an undigested mutant band (797 bp).

The *pax9^ci3038^* mutant line (NC_007128.7:g.38242475_38244047del) was generated through CRISPR/Cas9-mediated gene editing as described (Okeke et al., 2022). sgRNAs targeting sequences in the 5’UTR (5’-GCTAAACTGGACTCGGAAC-3’) and downstream of exon 2 (5’-GAAGAATTGGCCCAGCCAAG-3’) were designed, synthesized, and micro-injected at 100 ng/ml with 100 ng/ml Cas9 protein into one-cell stage Tubingen embryos. In the *pax9^ci3038^* isolate, a 1573-bp deletion removes the start codon, the Paired DNA-binding domain, and the conserved octapeptide. Translation from the next in-frame methionine would produce only short C-terminal fragments (55 or 14 amino acids in length for the long and short isoforms, respectively) lacking known functional domains. *pax9^ci3038^* fish were genotyped using primers F1 (5’-AGGAAACAGAAGGCGAATTTTC-3’) and R1 (5’-GGCTGCGGCCTTTACATGATA-3’), producing a 450-bp band for the deletion allele, and F3-WT (5’-AAAGCGATGTGGGATGATTC-3’) and R3-WT (5’-TATTCCGTTTCCACGTTTCC-3’), yielding a 392-bp wild-type band.

### Musculoskeletal staining

Combined Alcian blue and Alizarin red staining for cartilage and mineralized bone were performed on larvae and adults following standard acid-free procedures (Walker and Kimmel, 2007; Ullmann, 2011). Juvenile/adult fish were sorted according to size before staining and closely monitored during bleaching and trypsinization steps, which took 3 to 5 hours and up to 2 hours, respectively, depending on size. Alizarin red-only staining was performed on juvenile and adult fish using an ethylene glycol permeabilization method (Sakata-Haga et al., 2018). Live bone staining in larvae was performed by incubating fish in 0.0033% Alizarin red (139 μM, Acros Organics) in EM for 30 min at 28.5°C, followed by extensive washing prior to imaging under anesthesia. For sequential live bone staining, following Gonzalez Lopez et al. (2024), we first prepared separate stock solutions of 13 mM Alizarin red and 0.9 mg/ml Calcein green (Fisher) in 2% NaHCO_3_. These were then diluted 1:500 in system water and applied to fish from 7-9 dpf (Alizarin), then from 13-14 dpf (Calcein); the fish were then washed and imaged under anesthesia.

Muscles were stained with phalloidin. Larvae and juveniles were euthanized and fixed overnight in 4% paraformaldehyde in phosphate buffered saline (PBS) at 4°C, then repeatedly washed and permeabilized the following day in 1% Triton-X-100 in PBS. They were then stained overnight at 4°C in a 1:100 dilution of Alexa647-conjugated phalloidin (ThermoFisher) in PBS, washed in PBS, and imaged.

### Colorimetric and fluorescent in situ hybridizations

In situ probes used in this study include *alx1 and gata3* (Barske et al., 2018), *barx1* (Barske et al., 2016), *dlx2a* (Akimenko et al., 1994), *lhx6* (Paudel et al., 2022), *pax3a*, and *pax9*. Partial cDNAs for *pax3a* and *pax9* were amplified with the following primers: pax3a: 5’-ACCCCAAAACTACCCGAGAG-3’ and 5’-CTCGTGCCTCTGTGAGTTTG-3’; pax9: 5’-CTGGACTCGGAACAGGTCAG-3’ and 5’-CCGTTATTGATCGAATGCCCA-3’. Fragments were cloned into the pCR-Blunt II-TOPO vector (Life Technologies) and sequence-verified prior to plasmid linearization with EcoRV and BamHI, respectively, and in vitro transcription with Sp6 and T7 RNA polymerase (Roche). Colorimetric in situ hybridizations for *pax9* were performed on embryos and older larvae using the xylene permeabilization method (Vauti et al., 2020). Fluorescent in situ hybridizations were performed on embryos as previously described (Talbot et al., 2010). All embryos and larvae used for in situs were euthanized in 0.4% Tricaine then fixed in 4% paraformaldehyde overnight at 4°C or 3-4 hours at room temperature and stored in 100% methanol at −80°C until use.

### Immunostaining and nuclear labeling

Immunostaining was performed following Okeke et al. (2022) on 4% paraformaldehyde-fixed larvae using anti-HCS-1 (otoferlin, Developmental Studies Hybridoma Bank / NICHD, RRID:AB_10804296, 1:500) or anti-cleaved caspase 3 (CC3, Cell Signaling Technology 9661, RRID:AB_2341188, 1:500 for 4 dpf, 1:600 for 7 dpf), in combination with Alexa dye-conjugated secondary antibodies (1:300, Thermofisher). Nuclear staining to reveal general head morphology (after Sandell et al., 2018) was performed on fixed juveniles and larvae with DAPI (1:1000, Sigma D9564; overnight at 4°C) or DRAQ5 (1:1000, Abcam ab108410; 20 min at room temperature), respectively.

### Cell proliferation analysis

To assess whether cell proliferation in maxillary domain is altered in mutants, BrdU snapshot assays were performed as described (Barske et al., 2020), with minor modifications. Larvae (4 dpf) were incubated with 4.5 mg/ml BrdU in 15% dimethyl sulfoxide for 1-2 hours and then fixed in 4% PFA overnight at 4°C, passed through a methanol gradient, and stored at −80°C. Larvae were subsequently rehydrated, genotyped, digested with proteinase K (20 µg/ml) at 25°C for 30 min, postfixed in 4% PFA for 20 min, permeabilized in cold acetone at −20°C for 15 min, and then treated with 4 N HCl for 20 min at 25°C before immunostaining with anti-BrdU (1:200; Novus Biologicals NB500-169, RRID:AB_10002608) primary antibodies and Alexa568-conjugated secondary antibodies (1:300). The blocking solution contained 0.5% Triton X-100. DRAQ5 (1:300) was applied with the secondary antibody.

### Wnt activator treatment

To boost canonical Wnt signaling, embryos were treated with BIO (6-bromoindirubin-3’-oxime) (TOCRIS Bioscience), a small molecule inhibitor of glycogen synthase kinase-3 (Sato et al., 2004). BIO was dissolved in DMSO and stored in the dark at 4°C. Doses tested ranged from 0.5-5 μM (e.g. Nikaido et al., 2017), and were initiated at 1, 2, 3, 4, or 5 dpf as continuous or intermittent (every other day) treatments. If they survived treatment, larvae were euthanized at 8 dpf and processed for Alcian-Alizarin staining and imaging. The examples shown in Fig. 9F were treated with 0.5 μM BIO for 24 h from 4 to 5 dpf, then moved into fresh medium until 7 dpf, when they were placed back into medium containing the same concentration of drug until 8 dpf, when they were euthanized and processed for staining.

### Semi-quantitative reverse-transcriptase PCR (rt-PCR)

*pax9* transcript levels in el622 and ci3038 homozygous mutant fish were estimated by semi-quantitative rt-PCR. All-mutant clutches were generated by in-crossing adult mutants. Wild-type control samples were collected in parallel from the parental genetic background (Tübingen). Each biological replicate consisted of a pool of ten 24-hpf embryos that were frozen dry at −80°C. RNA was extracted from each pool using the RNAqueous-4PCR Total RNA Isolation Kit (Invitrogen), and 434.4 ng were used to synthesize cDNA with the High-Capacity cDNA Reverse Transcription Kit (Applied Biosystems). PCR was run on 2 μl cDNA using F1: 5’-GCCTCACTTAGAACCTGAAGC-3’ and R1: 5’-TGTGGGGAGAGAGAGCACT-3’ primers and the following cycling conditions: 95°C for 3 min, 35 cycles of 95°C for 15 s, 56 for 30 s, and 72°C for 30 s, then a final extension at 72°C for 5 min. Three biological replicates were run for each comparison, and *eef1g* levels were used for normalization (Okeke et al., 2022). Band intensities were measured with Image Lab (BioRad) and analyzed using Prism 10 (GraphPad).

### Imaging

Whole-mount skeletal preparations were imaged in 50% glycerol in 0.1% KOH on a Zeiss StereoDiscovery v8 microscope. Dissected skeletal preparations were mounted in 50% glycerol in 0.1% KOH and imaged on a ZEISS Axio Imager Z1. Colorimetric in situs were imaged on a Nikon NiE upright microscope. 3% methylcellulose was used to mount heads in order to acquire frontal views. Fluorescent in situs were imaged on a Nikon ECLIPSE Ti microscope. MicroCT was performed on one 2-year-old *pax9^el622^* mutant and one size-matched control at the Molecular Imaging Center at the University of Southern California (following Teng et al., 2018). For all live imaging analyses, larvae were anaesthetized with 0.4% Tricaine and carefully positioned in a drop of 0.2% agarose before imaging with a 20x or 40x objective on a Nikon ECLIPSE Ti microscope. Slow-motion movies were captured with a Samsung Galaxy S22 Ultra phone using the Super Slow-Mo feature with motion activation (1080p resolution, 960 fps, 32x slower playback speed). Individual clips of adult mutant (n=19) and control (n=13) fish were obtained; of these, the best-matched pair that illustrate divergent jaw protrusion behaviors were selected for inclusion as Movies S1 and S2.

### Data analysis

Osteoblasts, maxillary mesenchyme, proliferating, and Wnt-active cells were quantified in *pax9*; *sp7*:*EGFP*, *pax9; alx4a:DsRed*, *pax9*, and *pax9*; *TCF*:*EGFP* fish, respectively, using the Spot function in Imaris 9.80. As some datasets did not pass the Kolmogorov-Smirnov test for normality, we used non-parametric Mann-Whitney tests to evaluate differences between groups at p<0.05 in GraphPad Prism.

## RESULTS

### Midfacial retrusion in adult pax9 mutant zebrafish

To test whether Pax9 is functionally required in the cyprinid oral domain despite their evolutionary loss of teeth, we used TALENs to construct a new zebrafish mutant allele, *pax9^el622^*. This is a 10-bp deletion in exon 2 that causes a frameshift after amino acid 33 (a quarter through the conserved Paired DNA-binding domain) and premature termination of translation after 16 incorrect amino acids (Fig. 1A). *pax9^el622/el622^*mutants developed swim bladders and grew to adulthood at Mendelian ratios in the presence of wild-type or heterozygous siblings.

**Fig. 1.**
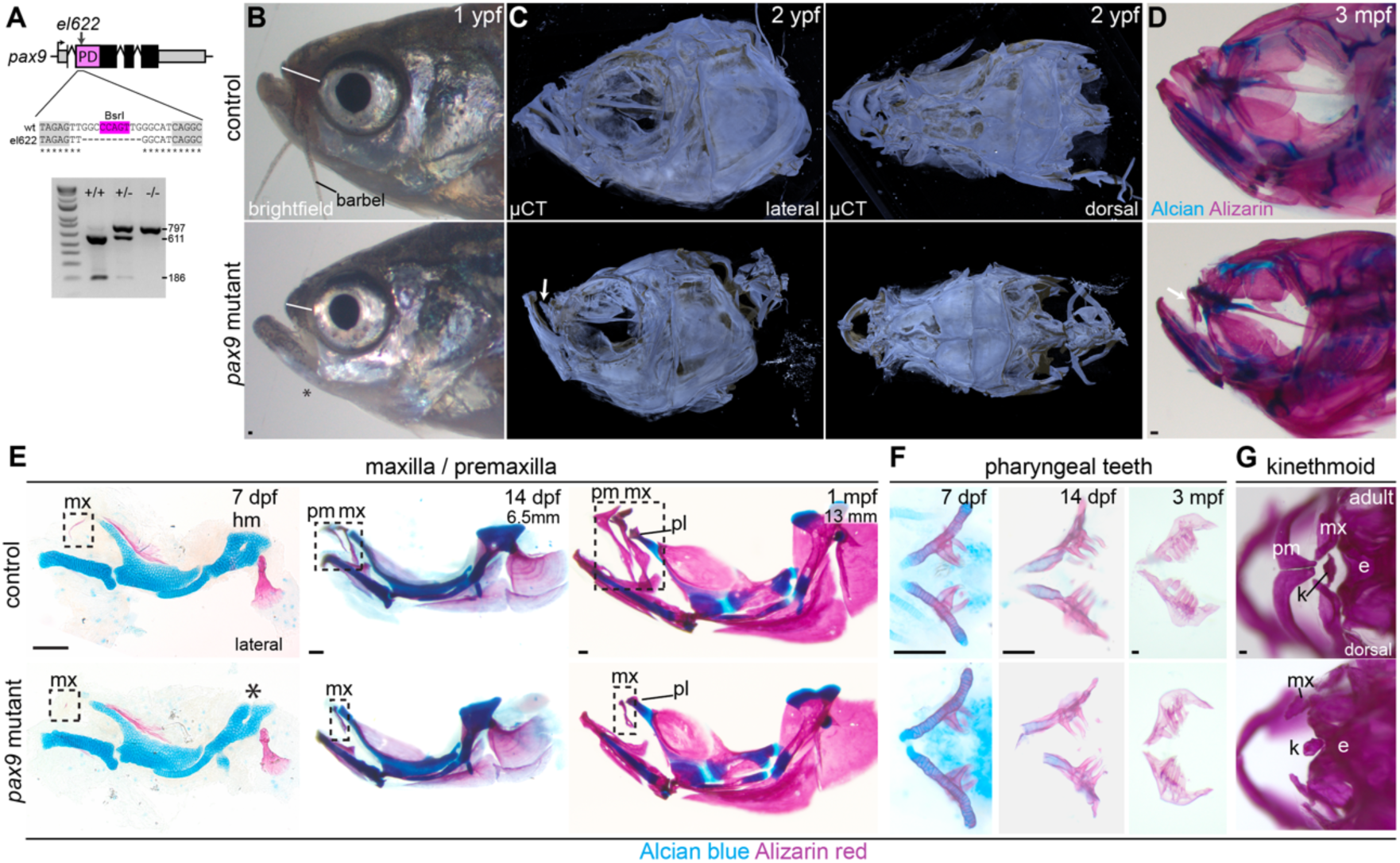
Developmental failure of upper jaw intramembranous bone formation in *pax9* mutant zebrafish. **A**, Schematic of *pax9^el622^*mutant allele. A 10-bp deletion in the paired domain (PD) removed a BsrI site, such that PCR amplification of the surrounding sequence and BsrI digest results in full cleavage of the product in wild-types and no cutting in mutants. **B**, Brightfield imaging of 1-year-old (ypf) fish reveals midfacial retrusion associated with overt shortening of the face anterior to the eye (white line) as well as missing barbels in mutants. **C**, Micro-CT of aged 2 ypf fish shows an even more pronounced underbite in mutants and apparent absence of most of the upper jaw complex. Both lateral and dorsal views are shown, n = 1 ea. **D**, Selective absence of the upper jaw bones is confirmed by Alcian Blue (cartilage) and Alizarin red (calcified bone) staining of younger adult mutants (3 mpf). White arrows in **C** and **D** point to the maxilla fragment. **E**, Developmental series of dissected skeletal preps between 7 dpf and 1 mpf shows that the maxilla (mx) initiates development on time, but is consistently shorter than in the control, failing reach full length both ventrally and dorsally relative to the palatine bone (pl) at 1 mpf. The premaxilla (pm) was never detected in mutants. A dorsal notch in the hyomandibular cartilage (asterisk) was occasionally noted in mutants. **F**, Dissected fifth ceratobranchials with attached pharyngeal teeth in larval, juvenile, and adults reveal no substantive differences in tooth number, position, or development in mutants. **G**, Dorsal view of representative normal and malformed kinethmoid bones (k) in control and mutant adult fish stained with Alizarin red. The mutant kinethmoid is nearly fused with the ethmoid (e). Scale bars = 100 μm.

Craniofacial dysmorphism externally resembling midfacial retrusion or mandibular prognathia became visually obvious in older mutant fish (Fig. 1B). Preliminary micro-CT imaging of a two-year-old mutant and sibling control revealed a striking absence of most of the upper jaw skeleton (Fig. 1C). To more closely examine this part of the skull, adult mutants and sibling controls were fixed and stained with Alizarin red for bone and Alcian blue for cartilage. Strikingly, mutants entirely lack the premaxilla bone as well as the ventral half of the maxilla (Fig. 1D), which normally articulates with the dentary bone of the lower jaw. These phenotypes were fully penetrant. Mutants also present a median fin anomaly that will be presented in a separate study. Pharyngeal teeth were indistinguishable in appearance and number in mutants compared with controls (Fig. 1F), consistent with a previous report that *pax9* expression is undetectable in these posterior tooth germs (Jackman et al., 2004). This supports that Pax9 is not unequivocally essential for tooth development.

To determine whether the upper jaw bones do not form at all in mutant fish or form but degrade during adult life, we performed an Alcian blue/Alizarin red time course of the jaw bones over the first month of life. The maxilla was first detected in mutants by weak Alcian and Alizarin staining at approximately 6-7 dpf, as in sibling controls (Fig. 1E), but failed to elongate after that. No mineralized premaxilla bones were observed in mutants at any stage. Mutants also occasionally presented a small dorsal notch in the hyomandibular cartilage (Fig. 1E, asterisk); this is reminiscent of but milder than the malformation reported in *pax9* morphants (Swartz et al., 2011).

Many teleost fish feed by protruding the premaxilla anteriorly to meet the mandible (Movie S1). In zebrafish, this is facilitated by the kinethmoid, an unpaired, mesoderm-derived sesamoid bone unique to cyprinids that is connected by ligaments to the neurocranium, premaxilla, maxilla, and palatine bones (Kague et al., 2012; Staab et al., 2012). Adult *pax9* mutants retain a deformed kinethmoid that in some cases is fused to the ethmoid bone, which separates the brain from the oral cavity (Fig. 1G). The facial muscle involved in premaxillary protrusion is the adductor mandibulae. The A1 division of this muscle originates on the preopercle and quadrate bones and inserts primarily on the lateral side of the dorsal half of the maxilla in wild-type adult fish (Hernandez et al., 2007; Diogo et al., 2008). This muscle is morphologically normal in *pax9* mutant fish at 14 and 28 dpf (11 mm standard length, SL) (Fig. 2A). The maxillary superficial tendon (MST) that connects the A1 to the maxilla (Tsai et al., 2023) also still forms in mutants, though appears less organized compared with sibling controls (visualized with the *scxa:*mCherry transgene at 7 and 20 dpf, Fig. 2B). This residual musculoskeletal connection appears sufficient to allow mutant fish to feed effectively in the laboratory environment, despite the lack of effective upper jaw protrusion (Movie S2). These results further suggest that the remnant of the maxilla that still forms in *pax9* mutants is the more dorsal component that bears the muscular insertion. Published live bone staining experiments in wild-type fish (Gonzalez Lopez et al., 2023) suggest that this dorsal part of the maxilla forms early, with the next phase of bone growth predominantly occurring at the ventral end, close to the corner of the mouth (also see Fig. 2C). Interestingly, in 8-dpf wild-types, the ventral half of this bone is ensheathed by *barx1^Gal4ff^; UAS:*nlsEGFP^+^ cells (Fig. 2D). These cells likely derive from the mesenchymal cell population under the oral epithelium previously shown to express both *barx1* and *pax9* at earlier stages (Swartz et al., 2011). Altered properties of these cells due to loss of *pax9* might therefore explain stalled elongation of the maxilla in mutants.

**Fig. 2.**
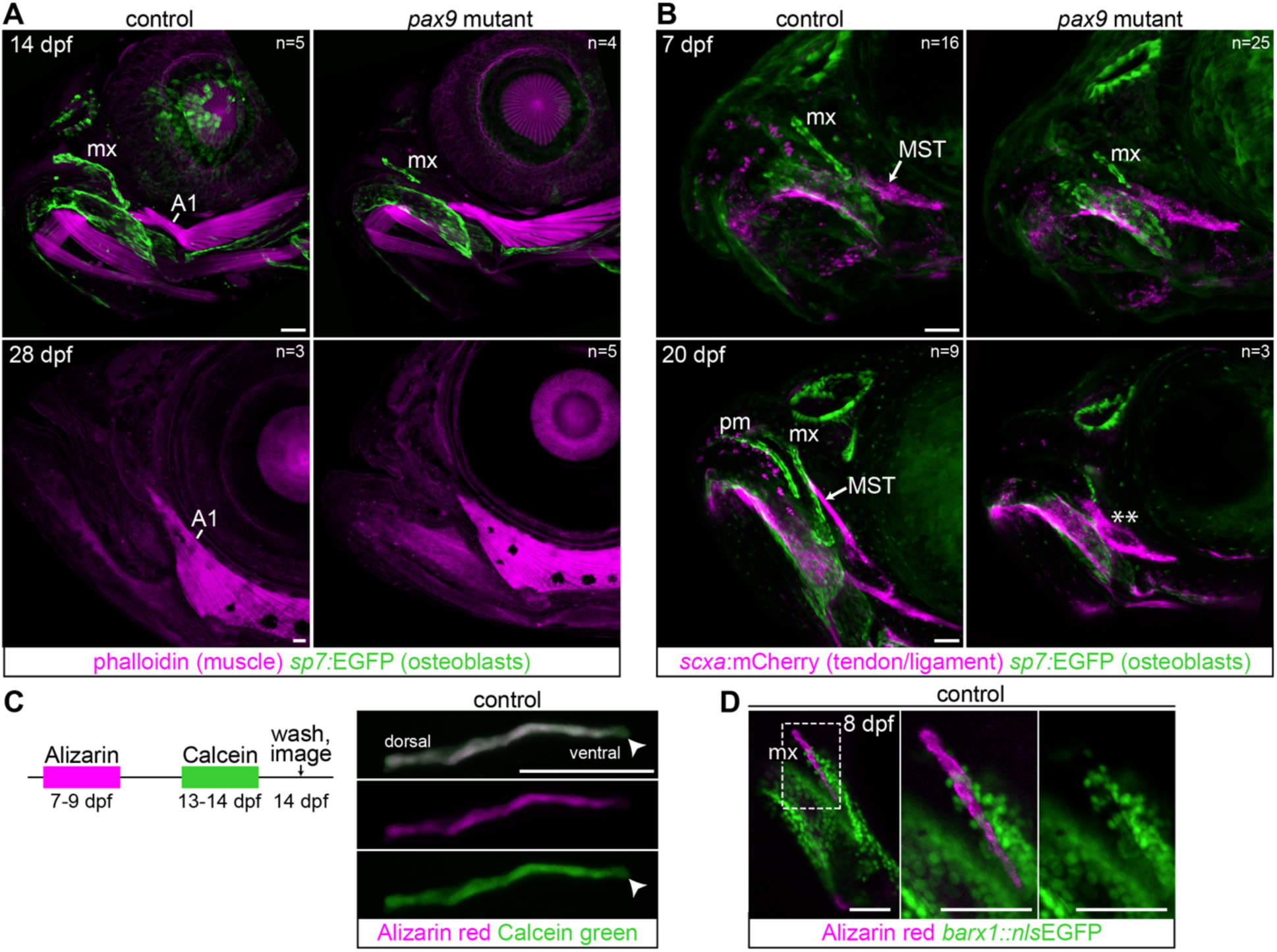
Retention of musculoskeletal attachments to the remnant upper jaw in *pax9* mutants. **A**, The A1 branch of the adductor mandibulae muscle involved in upper jaw mobility is not dysmorphic in *pax9* mutants at 14 or 28 dpf. Muscles are stained with phalloidin. Osteoblasts are labeled by *sp7:*EGFP in the top images. The maxilla (mx) is marked. **B**, The maxillary superficial tendon (MST) that connects the A1 to the maxilla is present in mutants at 7 and 20 dpf, but appears less organized relative to the control at the later stage (asterisks). Tendons and ligaments are marked by *scxa:*mCherry and osteoblasts by *sp7:*EGFP. N values reflect the total number imaged for that genotype/stage combination. **C**, Sequential bone staining by Alizarin red and Calcein green in wild-type fish shows that maxilla extends ventrally after 9 dpf (arrowhead). **D**, Live Alizarin staining of an 8-dpf *barx1^Gal4ff^; UAS:nlsEGFP* fish shows GFP^+^ cells closely ensheathing the ventral half of the maxilla. Scale bars = 50 μm.

### pax9 mutants lack sensory barbels

In addition to the missing upper jaw bones, adult *pax9* mutants also lack nasal and maxillary barbels (Fig. 1B). Barbels are sensory appendages flanking the mouth that are comprised of epidermis, goblet cells, pigment cells, taste buds, vasculature, sensory nerves, and fibroblasts, surrounding a central core of extracellular matrix (LeClair and Topczewski, 2009, 2010). These structures form during the juvenile period; in our facility, the first indications of barbel outgrowth appear around 1 mpf, at approximately 11 mm SL, and elongated structures are readily detectable at 16 mm SL (Fig. 3A-B, top panels). DAPI staining of juvenile *pax9* mutants at these stages show that these structures, too, entirely fail to form, and the soft tissues around the mouth are less convoluted compared with sibling controls (Fig. 3A-B, bottom panels). Lineage tracing confirmed previous reports that barbels contain mesenchyme (and nerves) of *so×10^+^* neural crest origin (Fig. 3C), ensheathed by ectodermal epithelial cells (LeClair and Topczewski, 2010; Mongera et al., 2013). By contrast, we find that placode-derived sensory neuromasts (Webb and Noden, 2015) are still present in the maxillary region of mutants, as visualized by HCS1 immunostaining of hair cells (Fig. 3E). This supports that the missing organs around the mouth are specifically later-developing structures built from cranial neural crest.

**Fig. 3.**
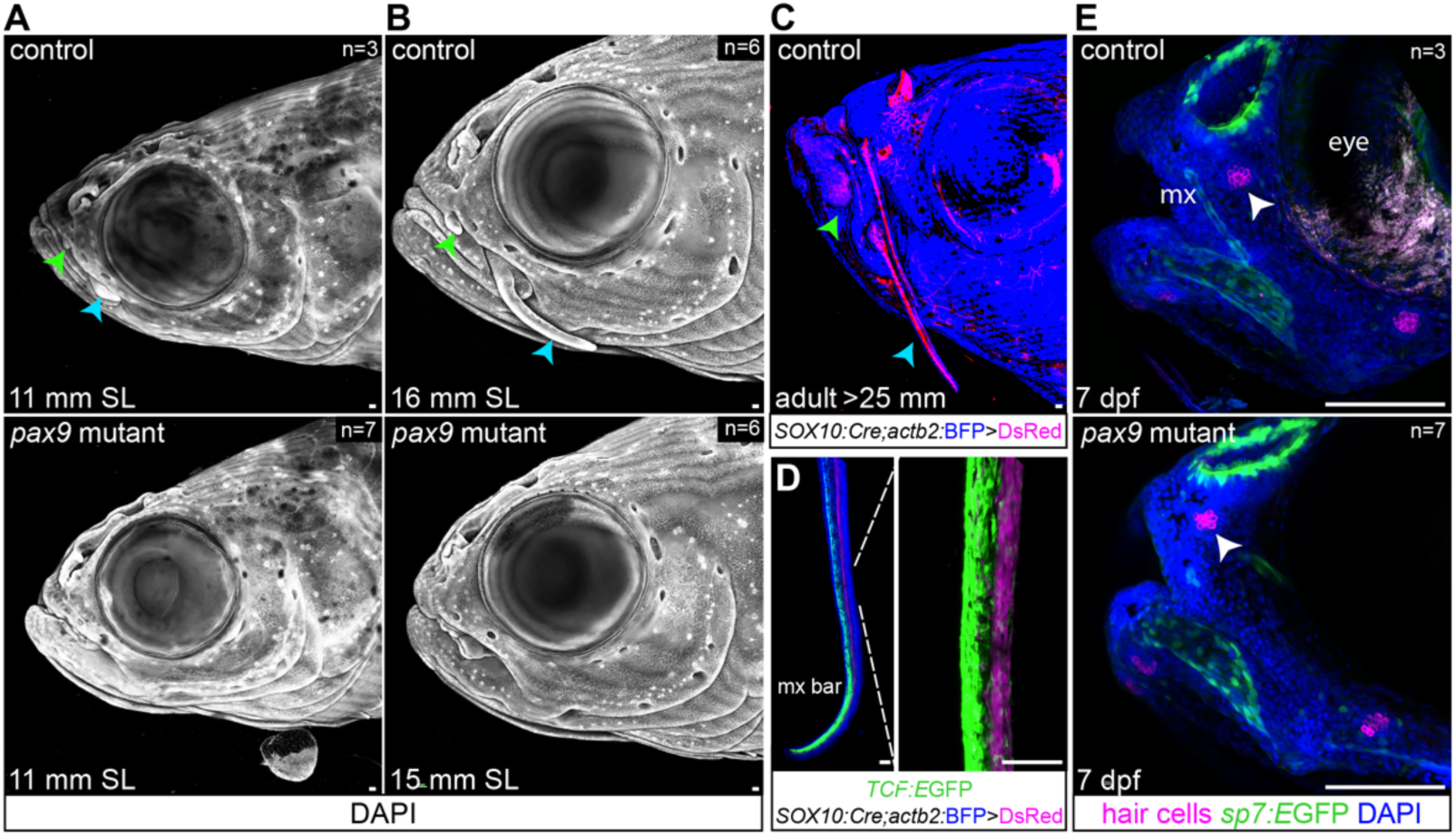
Pax9 is required for barbel formation. **A-B**, DAPI staining of juvenile specimens shows that maxillary (blue arrowheads) and nasal (green arrowheads) barbels bud and elongate between ∼11-16 mm in controls but do not appear in mutants. **C-D**, Lineage tracing of *SOX10*:Cre^+^ cells into adulthood confirms that barbels contain neural crest-derived cells, which also build the affected upper jaw bones. **D**, Barbels also contain a Wnt-responsive cell population (*TCF:*EGFP^+^) largely distinct from the neural crest-derived cells (DsRed^+^). **E**, The placode-derived neuromast (white arrowhead) positioned between the maxilla (mx) and the eye still forms in mutant fish. HCS1 immunostaining marks hair cells, *sp7:*EGFP labels osteoblasts and the nasal pit, and DAPI stains all nuclei. Images are maximum intensity projections. N values reflect the total number imaged for that genotype/stage combination. Scale bars = 100 μm.

### Severity of the pax9 mutant allele

Even though the juvenile and adult *pax9^el622^* skeletal phenotypes are so striking, the weakness of the larval craniofacial phenotype compared with the reported morpholino phenotype (Swartz et al., 2011) suggested that our allele might not be null. We therefore tested whether the mutant transcript was being degraded by nonsense-mediated decay and thus initiating transcriptional adaptation, a compensatory process that can ameliorate a mutant phenotype (Rossi et al., 2015; El-Brolosy et al., 2019; Ma et al., 2019). A slight, non-significant trend towards reduced transcription was observed by rt-PCR at 24 hpf (Fig. 4A), so some level of adaptation cannot be fully ruled out. We then used CRISPR/Cas9 to generate a larger deletion allele, *pax9^ci3038^*, in which a 1573-bp deletion removes most of the 5’UTR and exons 1 and 2 (Fig. 4B). rt-PCR for the 3’ end of the transcript revealed that enough 5’ initiating sequences remained to permit transcription of the mutant allele, again at similar levels as wild-type (Fig. 4C). *pax9^ci3038^* mutants present the same hyomandibula, barbel, maxilla, premaxilla, and kinethmoid phenotypes as *pax9^el622^*, with similarly normal palatal skeletons (Fig. 4D-G). This convergence in phenotype suggests that either both alleles are functionally null or both are compensated to the same minor degree.

**Fig. 4.**
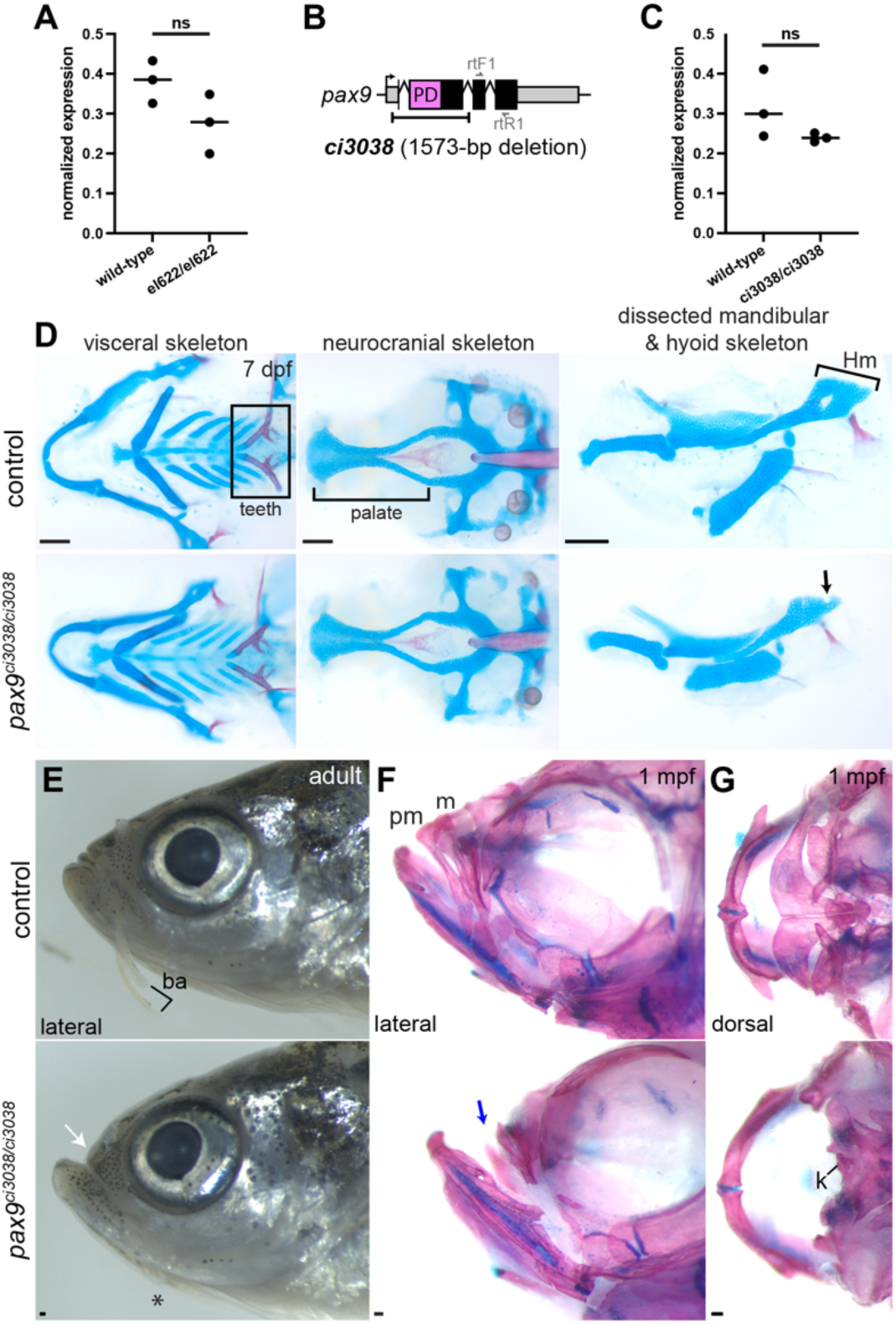
Second *pax9* deletion allele phenocopies the first. **A**, Semi-quantitative rt-PCR assays performed on 24 hpf samples reveal a non-significant (ns) decrease in transcription from the *pax9* el622 allele. Values calculated relative to *eef1g*. **B**, Schematic showing the approximate positions of the ci3038 lesion and the primers used for rt-PCR. **C**, rt-PCR at 24 hpf shows that the ci3038 allele is transcribed at levels not significantly different from wild-type. **D**, Homozygous *pax9^ci303/ci30388^* mutants have a mild hyomandibula phenotype (black arrow) and seemingly unaffected pharyngeal teeth and palate. **E-G**, Adult *pax9^ci303/ci30388^* mutants show midfacial retrusion (white arrow), lack of barbels (**E**), missing premaxilla and truncated maxilla (blue arrow, **F-G**), and malformed kinethmoid (k) (**G**). Specimens in **D**, **F-G** were stained with Alcian blue for cartilage and Alizarin red for bone. Scale bars = 100 μm.

### Oral pax9 expression persists after facial patterning stages

The unexpected reductions and losses of intramembranous bones and sensory appendages in zebrafish *pax9* mutants indicate broader, non-odontogenic requirements for Pax9 in the oral region. Lacking a robust fluorescent reporter for *pax9* expression, we in situ hybridizations to attempt to link these late-onset phenotypes to *pax9*^+^ cell populations. Prior studies focused on larval stages (up to 72 hpf) found strong mesenchymal expression at the lateral corners of the oral epithelium (both dorsal and ventral to the mouth), as well as a weaker central frontonasal domain (Jackman et al., 2004; Stock et al., 2006; Swartz et al., 2011), all of which we observe in 2-4 dpf samples (Fig. 5A). At 7 and 13 dpf, *pax9* expression is especially enriched in cells of the presumptive maxillary domain, clearly surrounding the maxilla itself in older specimens (Fig. 5B). Persistent expression in this region into early juvenile stages supports a role for Pax9 in morphogenesis and differentiation of late-developing facial structures like the maxilla.

**Fig. 5.**
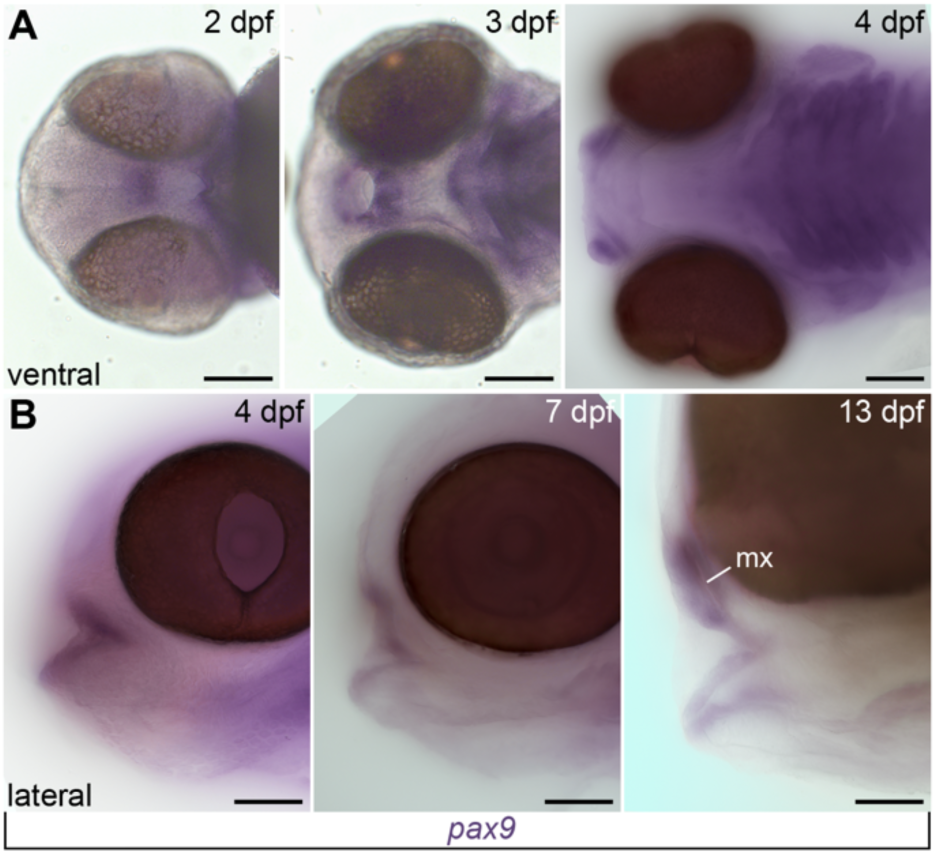
Oral *pax9* expression persists after facial patterning stages. Colorimetric in situs showing prolonged *pax9* expression around the mouth, weakly in parts of the oral epithelium and more strongly in the underlying mesenchyme. Clear mesenchymal expression around the maxilla is present in the 13 dpf specimen. **A**, ventral views; **B**, lateral views. Scale bars = 100 μm.

### No evidence of early patterning defect or mesenchymal cell loss

If Pax9’s requirement is at these later stages, earlier patterning of the frontonasal and peri-oral domains may be unaffected in mutants. By 36 hpf, cranial neural crest-derived mesenchymal cells have been patterned by signals from neighboring epithelia and transcription factor activity into distinct domains (Medeiros and Crump, 2012) (Fig. 6A). *dlx2a* marks crest-derived mesenchyme in the pharyngeal arches (Blentic et al., 2008) (Fig. 6C), including the *pax9^+^* condensed cells flanking the corners of the mouth (Fig. 6B) that co-express *lhx6* and *barx1* (Fig. 6D-E). *dlx2a*, *lhx6*, and *barx1* expression patterns in this region are not consistently different in mutants versus controls (Fig. 6C-E). Crest cells that migrate more anteriorly into the frontonasal domain turn on *pax3a*, *alx1*, and *gata3* instead (Zalc et al., 2015; Barske et al., 2018). These markers, too, are not consistently different between controls and mutants (Fig. 6F-H), indicating that the phenotype is likely not rooted in an early patterning defect.

**Fig. 6.**
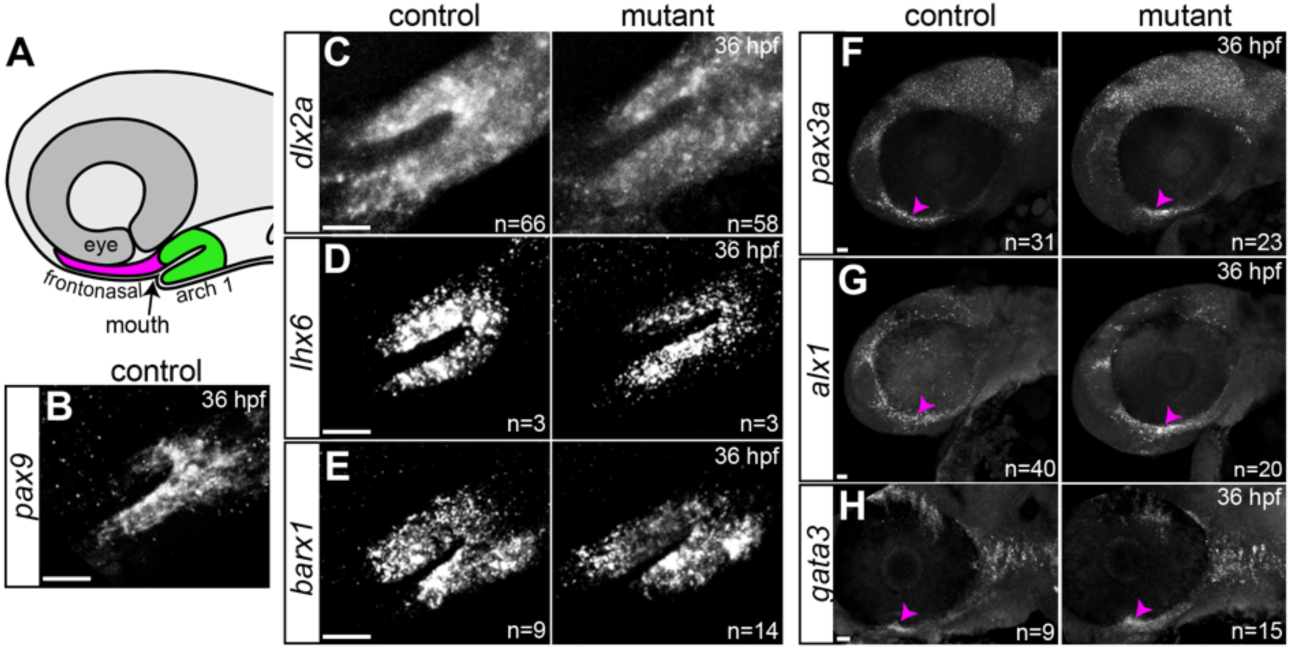
Normal expression of facial patterning genes in *pax9* mutants. **A**, Schematic of 36 hpf embryo head highlighting maxillary and mandibular cells of the first arch (green) that flank the oral epithelium and mouth opening and more anterior frontonasal cells (magenta). **B**, *pax9* is robustly expressed in mesenchymal cells around the mouth at 36 hpf. **C-E**, Mesenchymal expression of *dlx2a*, *lhx6*, and *barx1* in the same population of cells is not affected in *pax9^el622^* mutants. **F-H**, Frontonasal expression of *pax3a*, *alx1*, and *gata3* (magenta arrowheads) is not consistently different between mutants and controls. N values reflect the total number imaged for that genotype. Scale bars = 20 μm.

We next looked for other potentially aberrant features of mesenchymal cells in the maxillary region before the onset of bone and barbel development. At 3 dpf, the density and distribution of DRAQ5^+^ nuclei around the mouth were indistinguishable by eye between controls and mutants (Fig. 7A). Live imaging of a transgenic reporter for pre-skeletal mesenchyme (*fli1:*EGFP) likewise revealed no overt differences in mutants at 3 dpf (Fig. 7B). We then measured cell death and proliferation at slightly later stages by CC3 immunostaining and BrdU snapshot assays. No notable cell death was observed in the maxillary region of either controls or mutants at 4 or 7 dpf (Fig. 7C). The number of BrdU^+^ proliferating cells in the quadrant dorsal to the mouth was not different between mutants and controls at 4 dpf (Fig. 7D-E), though when we narrowed in to the region immediately dorsolateral to the mouth corner, where the maxilla will form, we detected a slight though significant decrease in mutants (p = 0.0178; Mann Whitney test, Fig. 7F). The mutants’ grossly normal appearance at these early stages supports that the defect occurs later during morphogenesis of the affected elements. This could be similar to how, in *Pax9* mutant mice, tooth development initiates in a seemingly normal jaw but then permanently stalls (Peters et al., 1998).

**Fig. 7.**
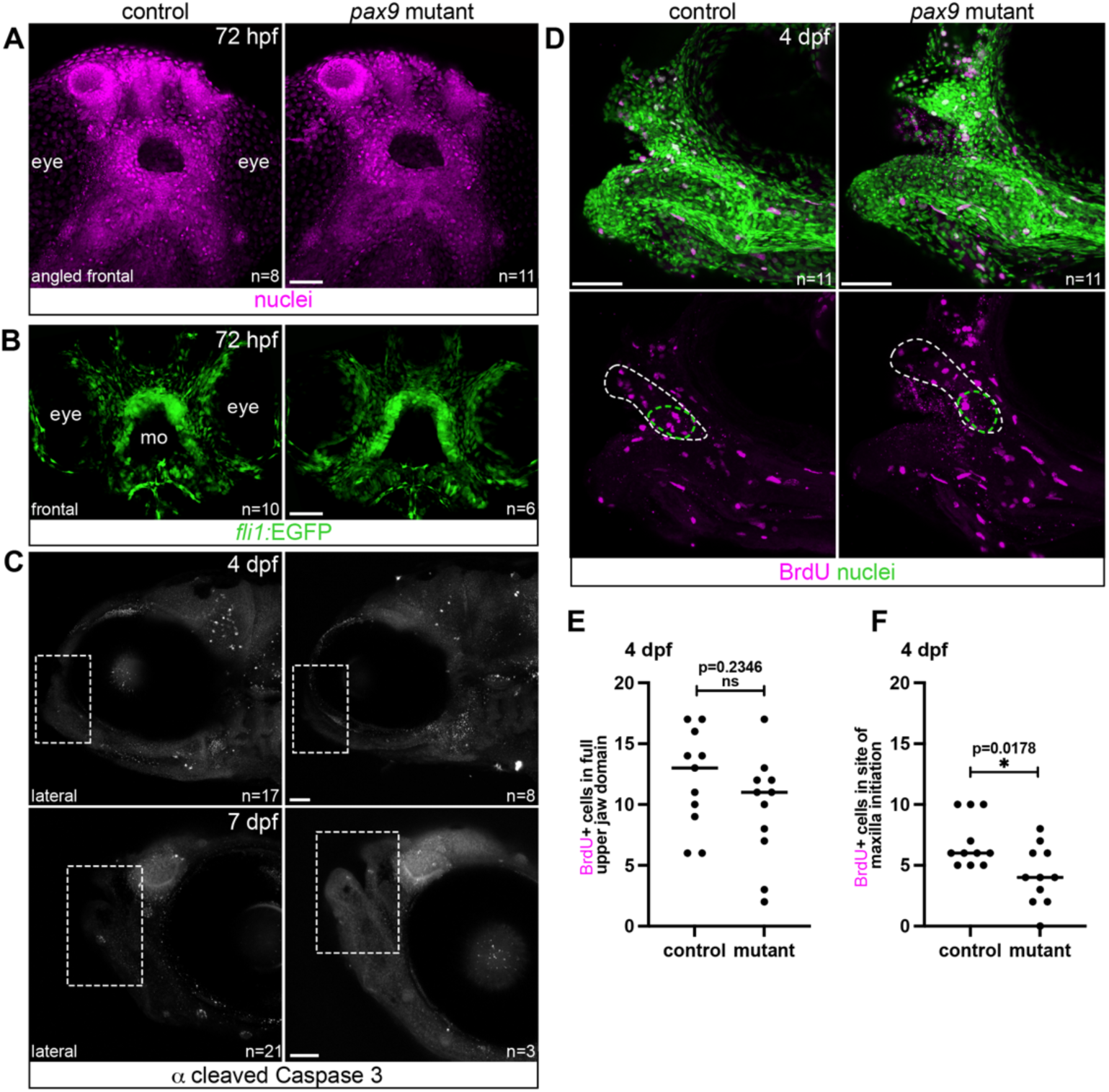
Oral mesenchyme is organized normally in larval *pax9* mutants. **A**, Staining of surface nuclei with DRAQ5 in 72 hpf control and mutant larvae reveals no overt abnormalities in the density or organization of cells around the oral opening (hole at the center of the image). **B**, Neural crest-derived mesenchyme marker *fli1:*EGFP is indistinguishable between controls and mutants at 72 hpf. mo, mouth. Angled frontal and frontal views are shown in **A** and **B**, respectively. **C**, Cleaved caspase 3 staining at 4 and 7 dpf revealed minimal cell death in both controls and mutants at 4 and 7 dpf. Outlined areas are the regions of interest around the mouth. Clear staining elsewhere in the head testifies to staining efficacy. **D-F**, BrdU snapshot assays at 4 dpf show similar levels of active proliferation in the full maxillary region (dashed white lines) in both controls and mutants (quantified in **E**), though fewer proliferating cells were counted in mutants in the more restricted domain in which the maxilla will initiate (green dashed line) (quantified in **F**; Mann Whitney non-parametric tests). Nuclei were stained with DRAQ5. N values in all panels are the numbers of imaged fish with the presented phenotype. Scale bars = 50 μm.

### Osteoblast deficiency in the maxillary region of pax9 mutants

Differences in mutants finally became apparent as maxillary osteoblasts began to differentiate and express *sp7:*EGFP (DeLaurier et al., 2010). EGFP^+^ cells sit directly adjacent to the mineralizing bone (Huycke et al., 2012) (Fig. 8A). At 7 dpf, we counted 12±0.8 EGFP^+^ osteoblasts around the forming maxilla in control larvae (mean±SEM; wild-type or heterozygous; n=8), but only 6.7±1.7 in mutants (n=9, p=0.0049, Mann Whitney test; Fig. 8B, D). One mutant in this batch had no EGFP^+^ cells at all. Significantly fewer EGFP^+^ cells in the maxilla were also observed at 14 dpf (30±2.8 in controls vs. 15±2.4 in mutants; n=7 and 6, respectively; p=0.0047, Mann Whitney test; Fig. 8C, E). EGFP^+^ cells in the position of the premaxilla became detectable in controls between 14 and 19 dpf depending on the overall growth of the larvae (e.g. Fig. 8C vs. H) but were never seen in mutants.

**Fig. 8.**
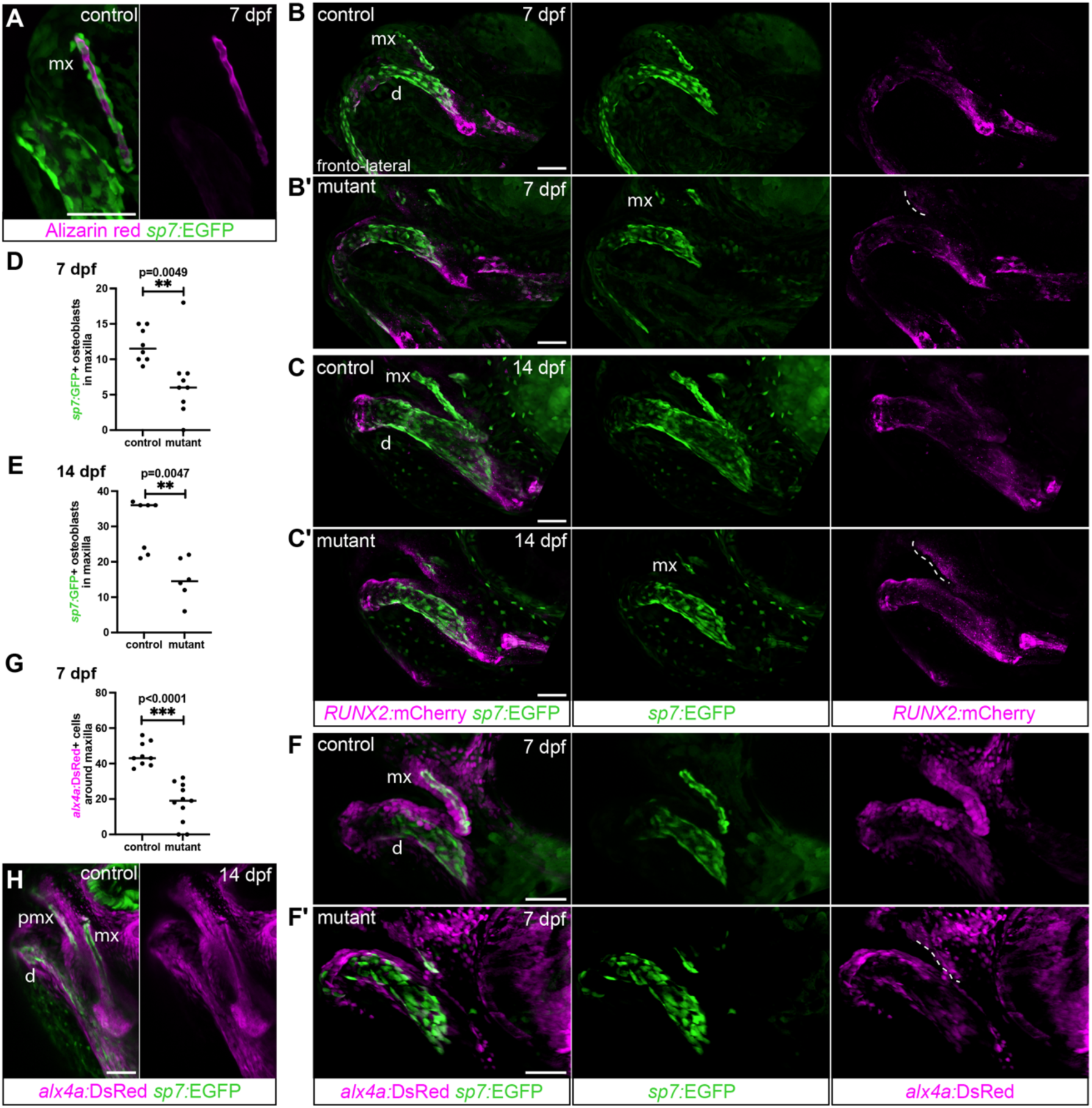
Reduction of the maxilla is associated with reduced osteoblast number. **A**, *sp7:*EGFP^+^ osteoblasts tightly encase the developing maxilla, stained with Alizarin red. **B-C**’, The number of *sp7:*EGFP^+^ osteoblasts in the maxilla region is significantly reduced in *pax9* mutants at 7 and 14 dpf. Cell counts compared by Mann Whitney tests in **D-E**. Weak granular *RUNX2*:mCherry expression reflective of osteoprogenitors is present in both *sp7:*EGFP^+^ osteoblasts and in the adjacent cell layer closer to the oral epithelium (dashed line). **F-F**’, *alx4a:*DsRed colocalizes with *sp7:*EGFP in osteoblasts around the maxilla and marks the immediate adjacent EGFP^-^ cell layers. Both EGFP^+^ and EGFP^-^ *alx4a:*DsRed^+^ cells are still detected in mutants though at reduced numbers (quantified in **G**, Mann Whitney test)). Dashed lines in **F**’ indicate EGFP^-^ domains of *alx4a:*DsRed closer to the mouth. **H**, By 14 dpf, the *alx4a:*DsRed^+^/*sp7*:EGFP^+^ premaxilla is also evident in a control fish. Images shown are maximum intensity (**A, C, H**) or 3D volume (**B, F**) projections, with Nikon DenoiseAI applied to the images in **C** for improved clarity. mx, maxilla; d, dentary; pmx, premaxilla. Scale bars = 50 μm.

We next used a reporter for *alx4a* (*alx4a:*DsRed (Nachtrab et al., 2013)), a gene known to be enriched in frontonasal neural crest-derived cells (Mitchell et al., 2021). DsRed was robustly expressed in maxillary and premaxillary *sp7:*EGFP^+^ osteoblasts of control larvae at 7 and 14 dpf, as well as in several surrounding mesenchymal cell layers (Fig. 8F,H). Again, fewer DsRed^+^ cells were associated with the maxilla in mutants at 7 dpf (45±2.2 in controls vs. 18±3.4 in mutants; n=9 and 11, respectively; p<0.0001, Mann Whitney test; Fig. 8F-G). Previous work in zebrafish has shown that in growing intramembranous bones, *sp7:*EGFP^+^ osteoblasts arise from initially EGFP^-^ cells recruited at the periphery of the ossification center (Huycke et al., 2012; Farmer et al., 2024). The *alx4a:*DsRed single-positive cells surrounding the double-positive domain may thus be poised to be recruited as osteoblasts, with this pool being deficient in the absence of *pax9*.

We also crossed in the *RUNX2*:mCherry reporter (Barske et al., 2020), which is expected to activate earlier in the osteoblast differentiation process than *sp7*:EGFP and then turn off as cells mature. At 7 and 14 dpf, we detect mCherry expression in the *sp7*:EGFP^+^ cells building the maxilla as well as in an adjacent anterior swath of cells under the oral epithelium (Fig. 8B-C). The mCherry fluorescence is fragmented rather than definitively cellular and cannot be confidently quantified; however, in mutants, we generally see a reduction in the domain immediately associated with the maxilla (overlapping the *sp7* osteoblast marker) but persistent expression closer to the oral epithelium. Strong mCherry expression under this epithelium is also noted in both mutants and controls as early as 3 dpf, before any osteoblast differentiation is apparent (Fig. 9A-B). The more centrally located mCherry^+^ cells lie within in the perichondrium of Meckel’s cartilage (lower jaw) and the ethmoid (palate), while the clusters at the corner of the mouth could be osteoprogenitors poised to be recruited to the maxilla (Fig. 9A, asterisks). We do not detect consistent differences between controls and mutants with this transgene at 3 dpf. Collectively, these observations suggest that the initial population of committed maxillary *sp7*:EGFP^+^ osteoblasts fails to expand in *pax9* mutants, potentially due there being few recruitable *alx4a:*DsRed^+^ cells, which may in turn derive from early *RUNX2:*mCherry^+^ osteoprogenitors around the mouth. We expect this differentiation failure occurs after 3 but before 7 dpf and may be confined to the maxillary subpopulation where we detected a small proliferation deficit at 4 dpf (Fig. 7F).

**Fig. 9.**
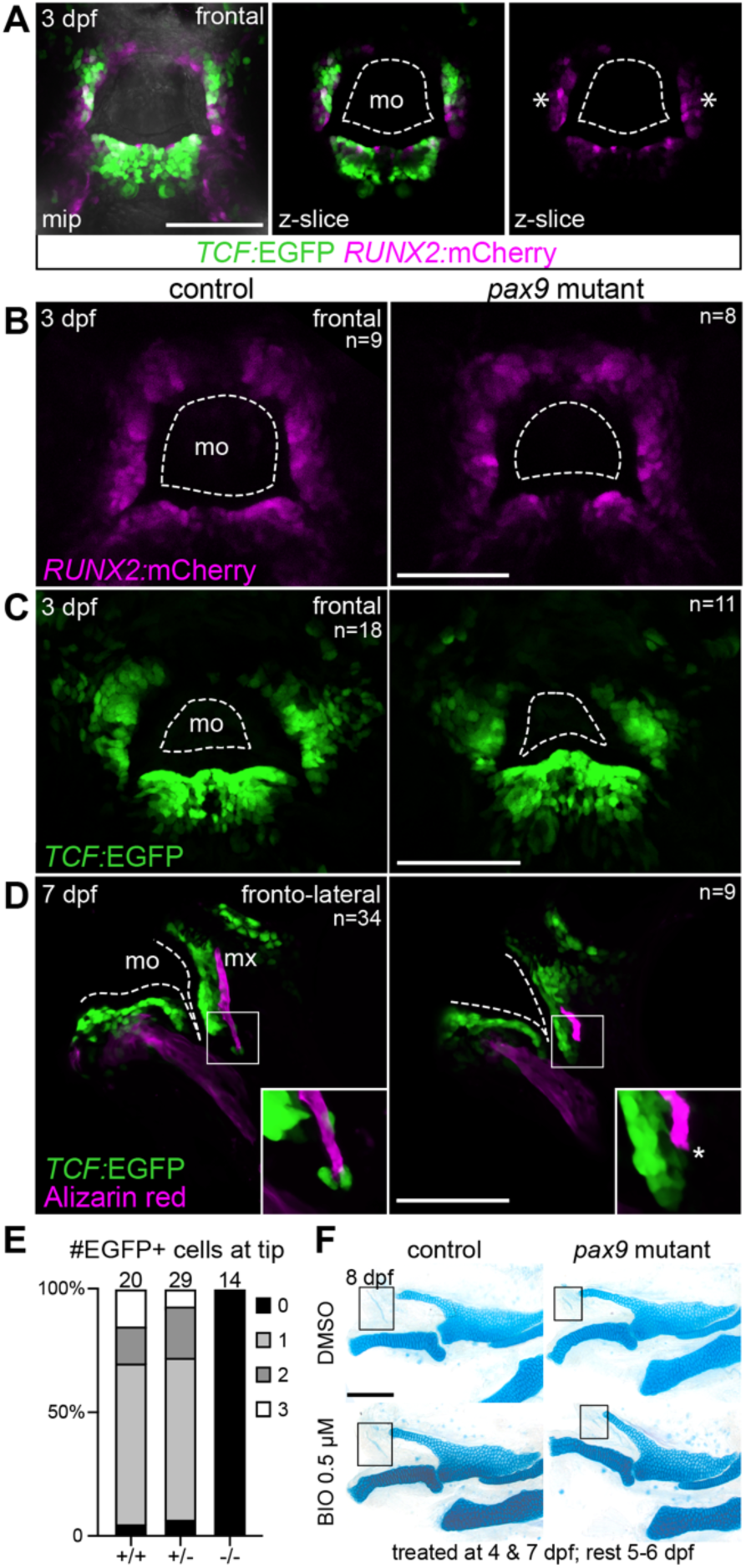
Oral Wnt signaling activity is not grossly altered by loss of Pax9 function. **A,** The Wnt reporter *TCF:*EGFP is strongly expressed in mesenchymal cells underlying the oral epithelium at 3 dpf, focused at the medial region of the forming lower jaw and the flanking dorsolateral domains. *RUNX2*:mCherry partially co-localizes, suggesting that some of the Wnt-active cells are osteoprogenitors. A maximum intensity projection is shown on the left, with z-slices on the right. Frontal views are shown; dashed line indicates the mouth opening (mo). **B-C**, Neither *RUNX2:*mCherry nor *TCF:*EGFP patterns in peri-oral mesenchyme are grossly different between control and mutant embryos at 3 dpf. **D**, Angled frontolateral views reveal persistent *TCF*:EGFP expression subjacent to the oral epithelium (dashed line) in control and mutant fish at 7 dpf. Fish were live-stained with Alizarin red before imaging. There is a gap between the peri-oral EGFP^+^ population and the maxilla in both controls and mutants. Images in **B-D** are maximum intensity projections. N values reflect the total number imaged for that genotype. **E**, Quantification of EGFP^+^ cells at the ventral end of the maxilla in wild-type, heterozygous, and mutant fish. Magnified views of these cells are shown in the insets in **D**. The total number for each genotype is listed at the top of the respective columns. **F**, Representative results from a BIO treatment showing no rescue of maxilla (boxed) size in mutants after boosting Wnt signaling. Scale bars = 100 μm.

### Bone and barbel phenotypes implicate Wnt signaling

One pathway that might work downstream of Pax9 in these oral-adjacent structures is canonical Wnt/β-catenin signaling. This pathway is essential for tooth progression (Chen et al., 2009) and intramembranous bone development (Day et al., 2005; Hill et al., 2005; Tran et al., 2010) and is also active during barbel outgrowth in zebrafish (Figueroa et al., 2015). In *Pax9* mutant mice, Wnt activity is reduced in the palatal shelves, due in part to expanded expression of Dkk inhibitors (Jia et al., 2017; Jia et al., 2020; Pina et al., 2024). This deficit contributes to their clefting phenotype, as four independent methods of reactivating Wnt – both genetic and pharmacological – were sufficient to rescue fusion of the secondary palate in mutants (Jia et al., 2017; Li et al., 2017; Jia et al., 2020). However, tooth development was not rescued in these models, indicating that Wnt is not the primary or only pathway dysregulated in *Pax9* mutant odontogenic tissue. To assess whether the loss of bone in our *pax9* mutant model might be associated with disturbed Wnt signaling, we crossed in the synthetic *TCF-Xla.Siam:GFP* Wnt reporter (hereafter *TCF:EGFP*) (Moro et al., 2012) and performed live imaging at 3 and 7 dpf. The oldest were live-stained with Alizarin red to label mineralized bone prior to imaging. Frontal views taken of control fish at 3 dpf reveal mesenchymal EGFP^+^ cells concentrated immediately under the oral epithelium (Fig. 9A,C), where we also observed strong *RUNX2:*mCherry (Fig. 9A-B). The ultimate fate of these double-positive cells is not yet known. EGFP-negative regions were noted at the midline dorsal to the mouth and at the lateral corners. In controls at 7 dpf, EGFP^+^ cells are still present around the mouth, filling much of the space between the oral epithelium and the forming maxilla (Fig. 9D). These Wnt-responsive cells do not ensheathe the bone as closely as the *sp7*:EGFP^+^ and *alx4a*:DsRed^+^ cells (Fig. 8A, F), except for 1-3 EGFP^+^ cells present at the ventral tip (insets in Fig. 9D), where the maxilla is actively growing (Fig. 2C). In mutants, the distribution, variability, and intensity of peri-oral EGFP expression was comparable to sibling controls at both stages. The only exception was that small population of EGFP^+^ cells around the ventral maxilla, which were never seen in mutants (Fig. 9D-E). Attempts to pharmacologically boost canonical Wnt signaling prior to the onset of the overt maxilla phenotype using the small molecule BIO (Sato et al., 2004) resulted in either no effect on skeletal development or lethality at the variety of doses and regimens tested (0.5-5 μm, initiating treatment at 1-5 dpf; representative results shown in Fig. 9F).

## DISCUSSION

The maxilla and premaxilla form relatively late in the larval period in zebrafish, past the limit of viability of many mutant lines, and little is therefore known about the cells and signals that control their development. The unexpected complete loss of premaxilla and stalled development of the maxilla observed in our *pax9* mutants expand the developmental requirements of this transcription factor to upper jaw bone development. Cubbage and Mabee (1996) noted that the premaxilla initiates development by the lateral edge of the maxillary head. It is therefore possible that the lack of premaxilla in *pax9* mutants could be a secondary consequence of the stalled growth of the maxilla. In other words, the maxilla may need to grow to a certain size or position to initiate adjacent condensation of the future premaxilla. Our further observation that mutants form apparently normal teeth deep in the pharynx but lack maxillary and nasal barbels motivate a model where Pax9 is not essential for tooth formation specifically, but instead is more generally required for building structures from condensed neural crest-derived mesenchyme around the mouth. This includes oral teeth, in species that have them. We accordingly hypothesize that null mutations in *pax9* in a non-cyprinid teleost species that retains teeth on the oral jaws would result in loss of these teeth as well as loss of the upper jaw bones and barbels.

Missing barbels have also been reported in zebrafish mutant for *ror2* (encoding a non-canonical Wnt receptor) (Dranow et al., 2023), *kctd15a/b* (encoding a potassium channel) (Heffer et al., 2017), *ccl33.2/ccl33.3* (encoding chemokines) (Zhou et al., 2018), and *duox* (encoding an enzyme required for thyroid hormone synthesis) (Chopra et al., 2019). Of these, we note that at least the *ror2* and *kctd15a/b* mutants also show dysmorphic and truncated maxillary and premaxillary bones, respectively (Heffer et al., 2017; Dranow et al., 2023). This convergence supports that the development of these bones and sensory structures is linked, potentially through their shared dependence on condensation of neural crest-derived mesenchyme in the oral region. Heffer et al. (2017) posited that the reduced body size of *kctd15a/b* mutant adults might be in part attributable to their loss of barbels, which carry taste receptors; i.e. they may have been eating less. Our anecdotal observation that *pax9* mutants are normal in size, despite having absent barbels and a much more dramatically malformed upper jaw, does not support this model.

We have not yet identified which downstream pathway disrupted by loss of *pax9* is responsible for the jaw phenotypes described here. Canonical Wnt reporter activity under the oral epithelium remains high in mutants (Fig. 9), and we were unable to find a dose or regimen of the Wnt activator BIO that had any impact on the maxilla phenotype. This result is consistent with pharmacological and genetic Wnt activation strategies not rescuing stalled tooth development in murine *Pax9* mutants, despite effectively reversing palatal clefting (Jia et al., 2017; Li et al., 2017). Further, if the *pax9* jaw phenotype had been due to broadly decreased Wnt activity, it might have manifested differently: in the mouse calvarium, loss of Wnt/β-catenin function causes progenitor cells to become chondrocytes instead of osteoblasts (Tran et al., 2010; Goodnough et al., 2012). We do not observe any ectopic cartilage in the jaw region of fish *pax9* mutants. We do note, however, that the small yet distinct group of *TCF:*EGFP^+^ Wnt-active cells coating the ventral tip of the maxilla in controls are missing in mutants (Fig. 9E-F). It has been proposed that zebrafish intramembranous bones grow by recruitment of cells in the surrounding mesenchyme to become osteoblasts, rather than only through intrinsic expansion of the initially specified population (Huycke et al., 2012; Teng et al., 2018; Dambroise et al., 2020; Farmer et al., 2024). We speculate that these Wnt-active cells might be newly recruited osteoblast progenitors contributing to ventrally-directed growth of the maxilla, a process that stalls out in mutants. Pax9 might therefore be required within mesenchymal progenitor cells to prime them for recruitment to the bone – in its absence, the cells may be blind to recruitment signals, which probably include Wnts (Gaur et al., 2005).

In contrast to our zebrafish mutant, the *Pax9* mutant mouse has a largely unaffected maxilla and premaxilla, only presenting mispositioned palatal processes of the maxilla associated with cleft secondary palate (Peters et al., 1998). We propose that lack of palatal clefting in our mutant fish is attributable to the zebrafish palate not requiring the extra step of palatal shelf elevation prior to midline fusion (Duncan et al., 2017) that occurs in mammals (Bush and Jiang, 2012). This is the point that the process goes awry in *Pax9* mutant mice (Zhou et al., 2013). We also consider the possibility that the maxilla and premaxilla bones might have different embryonic origins in fish and mammals. Recent lineage-tracing work in mouse has demonstrated that the osteoblasts that build the mammalian premaxilla derive from the *Alx3*^+^ frontonasal prominence, whereas those of the maxilla derive from the *Dlx1*^+^ maxillary prominence of the first pharyngeal arch (Iyyanar et al., 2023). No such definitive lineage-tracing has been performed for the zebrafish: the relatively late-forming maxilla and premaxilla were left out of early fate-mapping studies (Wada et al., 2005; Crump et al., 2006). However, we note that both bones form from *alx4a:*DsRed^+^ cells (Fig. 8F, H); this reporter clearly labels cells in the frontonasal domain, but whether it also turns on in cells derived from the distinct anterior *dlx2a*^+^ maxillary prominence has not yet been determined. In mammals, *Alx4* transcription does extend into the maxillary domain (Gray et al., 2004), so it is possible that the fish maxilla and premaxilla will prove to have the same respective origins in the first pharyngeal arch and frontonasal domains as mammals. Nevertheless, comparative embryology and paleontological work indicate that substantial rearrangement has occurred in this part of the face throughout vertebrate evolution, and that renaming of some of these bones may be in order. For example, one recent study used nerve mapping to highlight that the mammalian premaxilla bone may be more directly homologous to the septomaxilla than the premaxilla of non-mammalian tetrapods (Higashiyama et al., 2021). Unusual upper jaw features have also been noted in coelacanth and lungfish, two extant fishes of the sarcopterygian lobe-finned lineage that gave rise to tetrapods and eventually to mammals. Coelacanth have no maxilla and only a small premaxilla (DeLaurier, 2019), while the two bones are fused in lungfish (Janvier, 1996). The discrepant mouse and fish Pax9 mutant upper jaw phenotypes thus motivate additional work to clarify these bones’ developmental origins and distinct genetic requirements across vertebrates.

## ACKNOWLEDGEMENTS

We thank members of the Barske lab for helping with molecular biology experiments and imaging; Gage Crump for supporting establishment of the first *pax9* mutant line; Joshua Waxman and James Nichols for helpful discussions and for sharing the *TCF:EGFP* and *alx4a:DsRed* transgenic lines; the USC Molecular Imaging Center; Flynn Littleton and the CCHMC Division of Veterinary Services for fish care; and the CCHMC Bio-Imaging and Analysis Facility for microscope access and support. Funding for this project was provided to L.B. by the Cincinnati Children’s Research Foundation.

## Author contributions

Conceptualization, formal analysis, visualization: S.P. and L.B. Investigation: S.P., S.M., S.G., and L.B. Methodology: S.P. and L.B. Supervision and writing: L.B.

## Data availability

All data supporting these results are included in the manuscript and figures.

## Conflict of interest

The authors have nothing to disclose.

---

**Movie S1.**
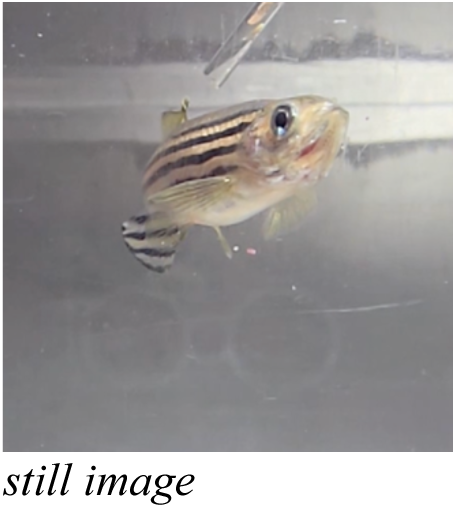
Wild-type adult zebrafish feeding behavior.

**Movie S2.**
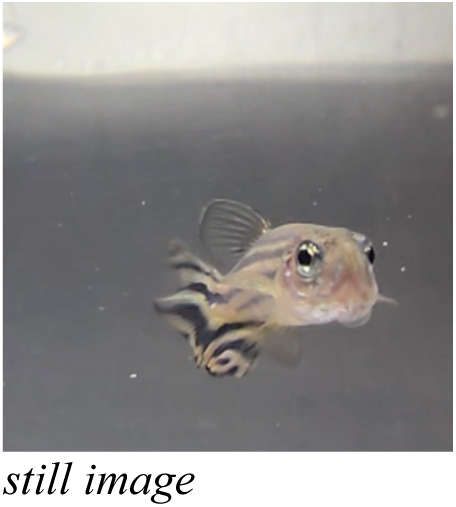
Adult *pax9* mutant zebrafish feeding behavior.

## Notes

### Competing Interest Statement

The authors have declared no competing interest.

